# Inter-unit variability and rapid depletion of 2,3-diphosphoglycerate in stored red blood cells: implications for transfusion practice in neurocritical care

**DOI:** 10.64898/2026.10.09.757834

**Authors:** Marta Peris, Juan Sahuquillo, Ana Ortuño, Veronica Pons, Marina Rierola, Maria Antonia Poca

**Affiliations:** Neurotraumatology and Neurosurgery Research Unit, Vall d’Hebron Institut de Recerca (VHIR), Vall d’Hebron Hospital Universitari, Vall d’Hebron Barcelona Hospital Campus, Passeig Vall d’Hebron 119-129, 08035 Barcelona, Spain; Independent Researcher and Scientific Advisor; Former Director, Neurotraumatology and Neurosurgery Research Unit, Vall d’Hebron Research Institute (VHIR), Barcelona, Spain; Department of Hematology, Vall d’Hebron Hospital Universitari, Vall d’Hebron Barcelona Hospital Campus, Passeig Vall d’Hebron 119-129, 08035 Barcelona, Spain; Blood and Tissue Bank, Vall d’Hebron Hospital Universitari, Vall d’Hebron Barcelona Hospital Campus, Passeig Vall d’Hebron 119-129, 08035 Barcelona, Spain; Department of Neurosurgery, Vall d’Hebron University Hospital, Passeig Vall d’Hebron 119-129, 08035 Barcelona, Spain; Department of Surgery, Universitat Autònoma de Barceloa (UAB), Bellatera, Spain

**Keywords:** Erythrocyte storage lesion, 2,3-Diphosphoglycerate, Liquid chromatography–mass spectrometry, Hemoglobin-oxygen dissociation, Blood transfusion

## Abstract

Red blood cell (RBC) transfusion aims to restore tissue oxygen delivery, yet a stored unit’s capacity to unload oxygen depends on 2,3-diphosphoglycerate (2,3-DPG), the principal allosteric modulator of hemoglobin–oxygen affinity. Randomized trials comparing fresher with older units have been consistently null, and all have used storage duration as the exposure variable. We asked whether storage age reflects the metabolic state of individual units. Twenty-four leukodepleted packed RBC units were sampled serially from day 3 to day 42 of standard cold storage. 2,3-DPG was quantified by liquid chromatography–mass spectrometry (LC–MS) with isotopically labelled internal standards, pH was measured in parallel, and trajectories were modelled by linear mixed effects with unit-level random intercepts. At day 3, 2,3-DPG spanned a twenty-fold range across units of identical age (0.8–14.7 µmol/g Hb; median 4.8): some were already near-depleted at the earliest sampled timepoint, while others retained near-physiological concentrations. This heterogeneity collapsed rapidly - median values fell to 27% of day-3 levels by day 7 and by more than 90% by day 14 (β = −0.298 µmol/g Hb per day, p < 0.001) - and were below the limit of detection in most units after day 17. pH declined monotonically to 6.40 ± 0.04 by day 42 (β = −0.0163 units per day, p < 0.001). Units of identical storage age therefore differ markedly in the metabolite that governs oxygen unloading, and virtually all are depleted long before the 42-day expiry, indicating that storage-age dichotomies misclassify the exposure they are intended to represent.

## Introduction

The primary objective of blood transfusion is to maintain or restore adequate oxygen delivery to tissues. However, prolonged storage under blood bank conditions leads to the development of the so-called “storage lesion”, a multifactorial process characterized by progressive biochemical, metabolic, and structural deterioration of red blood cells (RBCs). These changes include progressive metabolic depletion, disturbances in acid–base homeostasis, and increasing cytoskeletal instability [1,2]. During storage, the continued reliance of RBCs on glycolysis leads to gradual acidification of the suspension medium, a process widely recognized as a central mediator of storage-associated metabolic derangement and membrane remodeling [1,3,4]. This acidic milieu compromises cytoskeletal integrity, resulting in increased osmotic fragility (OF) and a higher rate of time-dependent hemolysis [1,3]. Consistent with this, Peris et al. recently demonstrated that OF increases significantly after 28 days of storage when RBCs are exposed to hypotonic stress, reflecting progressive impairment of membrane stability [3].

2,3-Diphosphoglycerate (2,3-DPG) is produced via the Rapoport–Luebering shunt, a glycolytic side pathway largely restricted to mammalian RBCs. Acting as a potent allosteric effector of hemoglobin (Hb), 2,3-DPG binds reversibly to deoxygenated Hb, reduces its oxygen affinity, and thereby promotes oxygen release to peripheral tissues [5–7]. Under standard blood bank conditions, 2,3-DPG concentration declines in a predictable manner, reaching very low levels after about two weeks and becoming nearly exhausted by 21 days of storage [7,8]. The loss of 2,3-DPG induces a left-shift in the oxyhemoglobin dissociation curve and reduces P_50_— the partial pressure of O_2_ at which hemoglobin is 50% saturated—thereby impairing the unloading of oxygen to tissues. Neurocritical patients commonly experience impaired cerebral oxygen delivery, rendering brain tissue highly sensitive to small shifts in hemoglobin–oxygen affinity. According to the Siggaard-Andersen classification, “high-affinity hypoxia” is a form of tissue hypoxia in which arterial oxygen tension and Hb concentration are normal, but Hb-O_2_ affinity is abnormally increased (reduced P_50_), thereby limiting oxygen unloading and tissue extraction despite adequate arterial oxygen content [9–11].

Our study aims to re-examine the temporal trajectory of 2,3-DPG depletion in stored RBC units using Liquid Chromatography–Mass Spectrometry (LC–MS). Unlike traditional enzymatic assays, LC–MS offers superior analytical specificity and sensitivity, enabling accurate quantification of phosphorylated glycolytic intermediates such as 2,3-DPG, particularly at the low concentrations encountered during prolonged storage [12–14]. By re-examining prior enzymatic assay–based findings using a more specific analytical approach, our objective was to elucidate the mechanisms underlying storage-dependent declines in 2,3-DPG and to assess the clinical relevance of these changes for the transfusion performance of RBCs in neurocritical patients, thereby contributing to the ongoing debate regarding the risk–benefit balance of RBC transfusion in acute brain injury.

## Study design and methods

This work is part of a broader research program conducted by the Neurosurgery and Neurotraumatology Research Unit at Vall d’Hebron University Hospital, dedicated to advancing strategies that optimize oxygen transport in neurocritical care. Ethical approval for the full project was granted by the Ethics Committee of the Vall d’Hebron Research Institute (protocol ID: PR-AG-411/2017). All procedures were carried out in accordance with Spanish law and the relevant European Union directives.

### 2.1. Leukodepleted packed red blood cell concentrates (pRBC)

Twenty-four leukodepleted pRBC units were obtained from our institution’s Blood and Tissue Bank, following all regulatory requirements, including adherence to EU Regulation 2016/679 governing personal data protection. Approximately 450 mL of whole blood from healthy volunteer donors was collected in CPD anticoagulant. Following centrifugation, leukocytes and plasma were removed, and the resulting RBC fraction was resuspended in 100 mL of SAGM (Saline, Adenine, Glucose, and Mannitol) storage solution. The pRBC units were delivered to the research team 3 days after donation, maintained at 1–6 °C, and examined from storage day 3 through day 42.

### 2.2. Liquid chromatography–mass spectrometry

For sample preparation, 250 μL of pRBC concentrate was obtained using heparinized collection tubes and then mixed with 500 μL of 1 M phosphate-buffered saline (PBS) and 30 μL of pure perchloric acid (HClO4) prior to analysis. Centrifugation at 12,000 rpm for 5 minutes at 4 °C produced a brown pellet and a clear, slightly light-brown supernatant. A 500 μL aliquot of the supernatant was then neutralized with 56 μL of 2.5 M potassium carbonate (K_2_CO_3_), generating a characteristic white precipitate. The mixture was then centrifuged again at 4 °C and 5,000 rpm for 10 minutes, and 400 μL of the final supernatant were collected and stored at −80 °C, the optimal temperature for metabolite preservation [14]. Preprocessed samples were maintained on dry ice during transport to the Applied Metabolomics Laboratory at the Instituto Hospital del Mar de Investigaciones Médicas (IMIM), where analyses were performed.

2,3-DPG was analyzed by LC–MS using a Waters ACQUITY UPLC system (Waters Corporation, Milford, MA, USA) coupled to a Waters Xevo TQS Micro triple-quadrupole mass spectrometer (Waters Corporation, Milford, MA, USA) equipped with an electrospray ionization (ESI) source. Quantification was performed by external calibration with defined analyte concentrations, complemented by isotopically labeled internal standards (^13^C-substituted analogs) that were added to both calibration solutions and sample matrices. Metabolite levels were determined by comparing the analyte and internal-standard peak areas. 2,3-DPG concentrations were reported relative to hemoglobin (µmol/g Hb), as this unit best reflects the physiologically relevant interaction between 2,3-DPG and hemoglobin and minimizes variability due to hematocrit [15]. Expressing the value per gram of Hb also normalizes for variations in hematocrit or red-cell mass between samples, thereby permitting direct comparison of metabolic status independent of cell count.

### 2.3. Potential of hydrogen (pH)

The measurement of pH was carried out using a specialized blood sample pH meter (InLab Micro Pro-ISM, Mettler-Toledo, Spain). The blood units were stored under the same conditions as the blood bank, at 4°C. However, pH measurements were taken at room temperature (20-23°C) to determine the pH of the bags under the conditions of routine blood transfusions.

Ensuring that the measurement temperature was very similar between measurements was considered essential, as it is a factor that influences the final pH value.

### 2.4. Statistical analysis

All statistical analyses were performed using R (v 4.5.2) within RStudio (v 2026.01.2). Descriptive statistics were calculated for all variables. Distributional properties were evaluated using exploratory graphical methods, including histograms and Q–Q plots, with particular emphasis on the normality of model residuals, consistent with the assumptions of linear mixed-effects modelling. Variables showing approximately symmetric distributions are presented as mean ± standard deviation (SD), whereas skewed variables are summarized as median and range. <u>Exploratory analysis</u>: Individual-level trajectories of 2,3-DPG were visualized using spaghetti plots to characterize within-unit decline patterns, and boxplots were used to summarize the distribution of 2,3-DPG and pH at each storage day. All figures were generated in R using the *ggplot2* package. These descriptive visualizations guided subsequent model specification. <u>Rationale for Mixed-Effects Modeling</u>: Because both 2,3-DPG and pH were measured repeatedly within the same pRBC units—and the number of observations varied across units—the longitudinal data were analyzed using linear mixed-effects models (LMMs). Storage time and pH were highly collinear, as expected from glycolytic metabolism. Including both predictors simultaneously would destabilize fixed-effect estimates. Therefore, two independent LMMs were fitted: 1. **Model 1:** 2,3-DPG ∼ Storage day + (1 | Unit) and **Model 2:** 2,3-DPG ∼ pH + (1 | Unit) Both models included random intercepts for each pRBC unit. Model assumptions (linearity, normality of residuals, and homoscedasticity) were assessed via diagnostic plots. Fixed-Effect Testing and Inference: Fixed effects were evaluated using Wald t-tests with Satterthwaite degrees of freedom, as implemented in *lmerTest*. Statistical significance was defined as *p* < 0.05. Model-based contrasts and estimated marginal means were obtained using the *emmeans* package when relevant [16]. <u>Data Management</u>. The study adhered to FAIR data principles. The complete anonymized dataset, metadata, and variable dictionary will be made available through Zenodo (DOI provided in final submission).

## Results

### 3.1. 2,3-DPG Dynamics During Storage

2,3-DPG concentrations declined rapidly and consistently across all leukodepleted packed RBC units. At the Day 3 baseline, median 2,3-DPG was 4.8 µmol/g Hb (range 0.8–14.7 µmol/g Hb), indicating substantial inter-unit variability early in storage. Despite this wide dispersion at baseline, all units exhibited a marked early decline in 2,3-DPG levels. By Day 7, median concentrations decreased to 1.3 µmol/g Hb (range 0.2–5.4), corresponding to approximately 27% of Day 3 levels (**Figures 1–2**). Beyond Day 7, 2,3-DPG concentrations continued to diminish; by Day 10, median values fell to 0.58 µmol/g Hb, and by Day 14 to 0.36 µmol/g Hb, representing a >90% reduction from baseline. After Day 17, most samples were below the LC– MS detection limit. Spaghetti plots revealed heterogeneous baseline concentrations but a uniform downward trajectory across all units, with the steepest decline occurring between Days 3 and 7 (**Figure 2**). <u>Linear Mixed-Effects Modeling</u>: A mixed-effects model with random intercepts demonstrated a strong association between storage duration and declining 2,3-DPG levels. Storage day was significantly associated with lower 2,3-DPG (β = −0.298 µmol/g Hb per day, robust SE = 0.053; t(18.1) = −5.59; *p* < 0.001). This corresponds to an estimated weekly decline of ≈2.1 µmol/g Hb. Model-based contrasts estimated a decrease of −1.19 µmol/g Hb from Day 3 to Day 7 (95% CI −1.64 to −0.75; *p* < 0.001) and −4.17 µmol/g Hb from Day 3 to Day 17. The model-supported pattern aligns with descriptive statistics, confirming rapid early depletion followed by near-complete exhaustion by Day 17.

**Figure 1.**
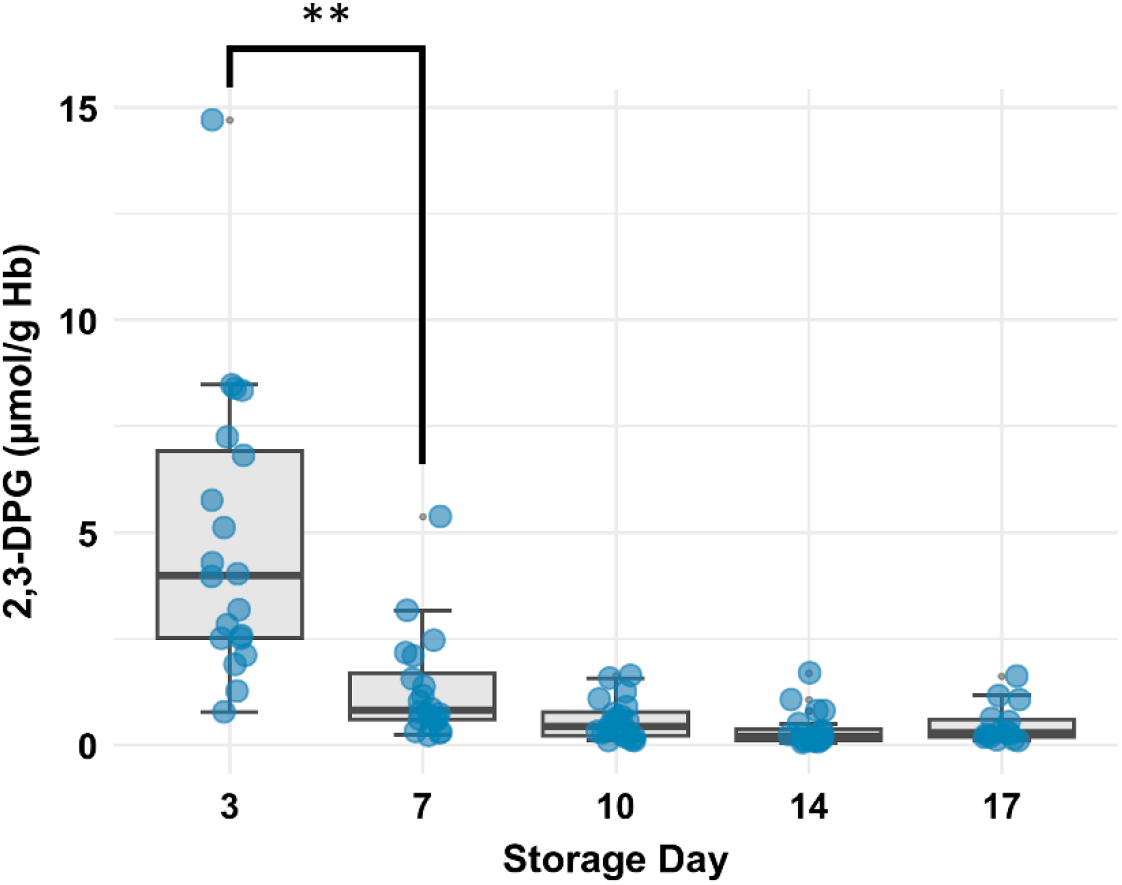
Changes in 2,3-DPG concentration during RBC storage. Box plots show 2,3-DPG concentrations (µmol/g Hb) from day 3 post-extraction through day 17 of storage. After day 17, 2,3-DPG levels became undetectable.

**Figure 2.**
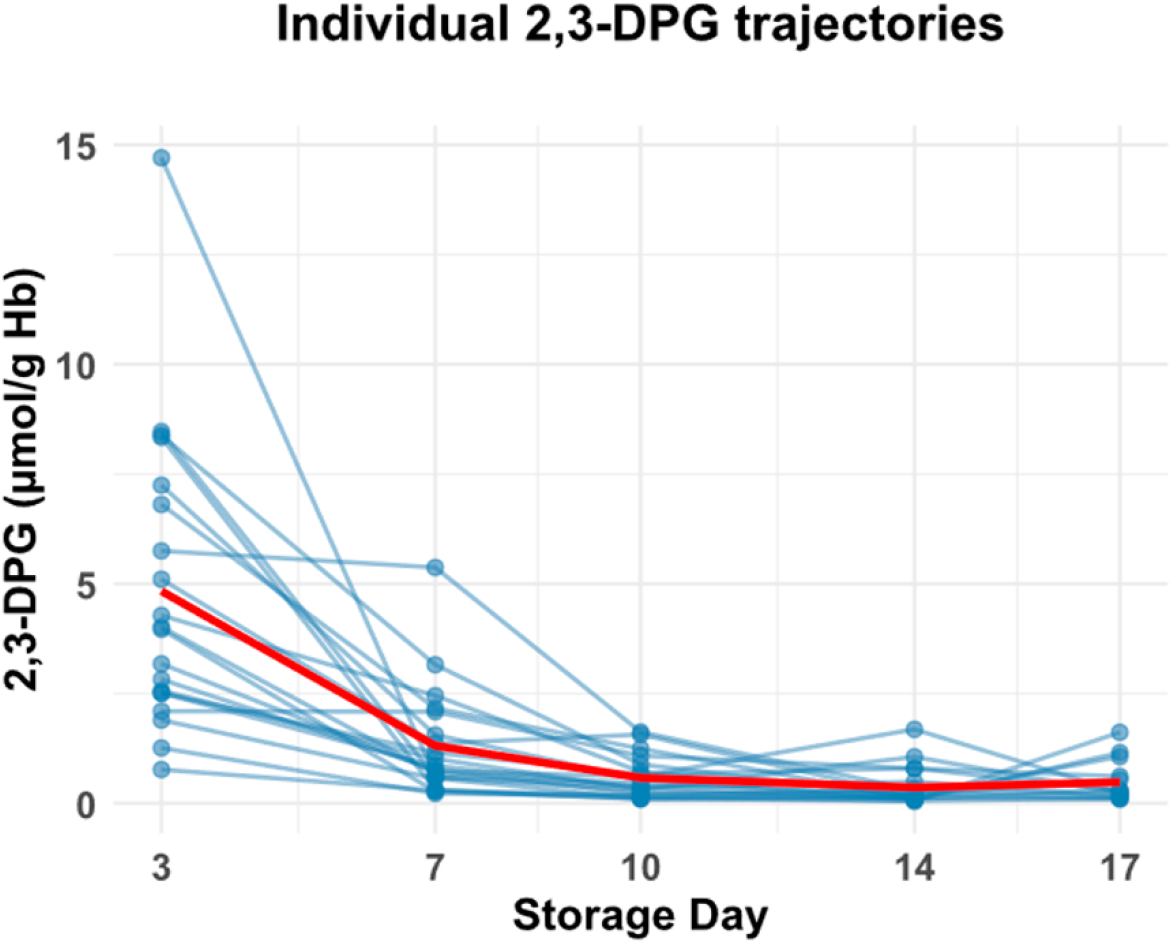
Spaghetti plot displaying individual 2,3-DPG trajectories (thin blue lines) for each leukoreduced pRBC unit. Although baseline concentrations vary substantially across bags, all units show a rapid early decline, with the steepest reduction occurring between Day 3 and Day 7. The population-level mean trajectory (thick red line) highlights the uniform, progressive loss of 2,3-DPG over time, culminating in near-complete depletion by Days 14–17. The plot also emphasizes the wide inter-bag variability in Day 3 post-extraction 2,3-DPG levels, despite the consistent downward trend thereafter.

### 3.2. Evolution of pH During Storage

The dataset included pH measurements from 24 pRBC units across 11 storage days (Days 3–42). On Day 3, the mean pH was 7.06 ± 0.06 (range 6.96–7.17). A progressive and nearly monotonic decline was observed thereafter, with mean pH reaching 6.40 ± 0.04 (range 6.31–6.46) by Day 42. Boxplots with individual data points (**Figure 3**) showed modest inter-unit variability at early time points and marked convergence toward more acidic values at later days. A linear mixed-effects model with a random intercept for *bag_id* demonstrated a strong association between increasing storage duration and decreasing pH (β = −0.0163 pH units/day; SE = 0.00033; t(236) = −49.6; p < 0.001). This corresponds to an average decline of approximately 0.11 pH units per week.

**Figure 3.**
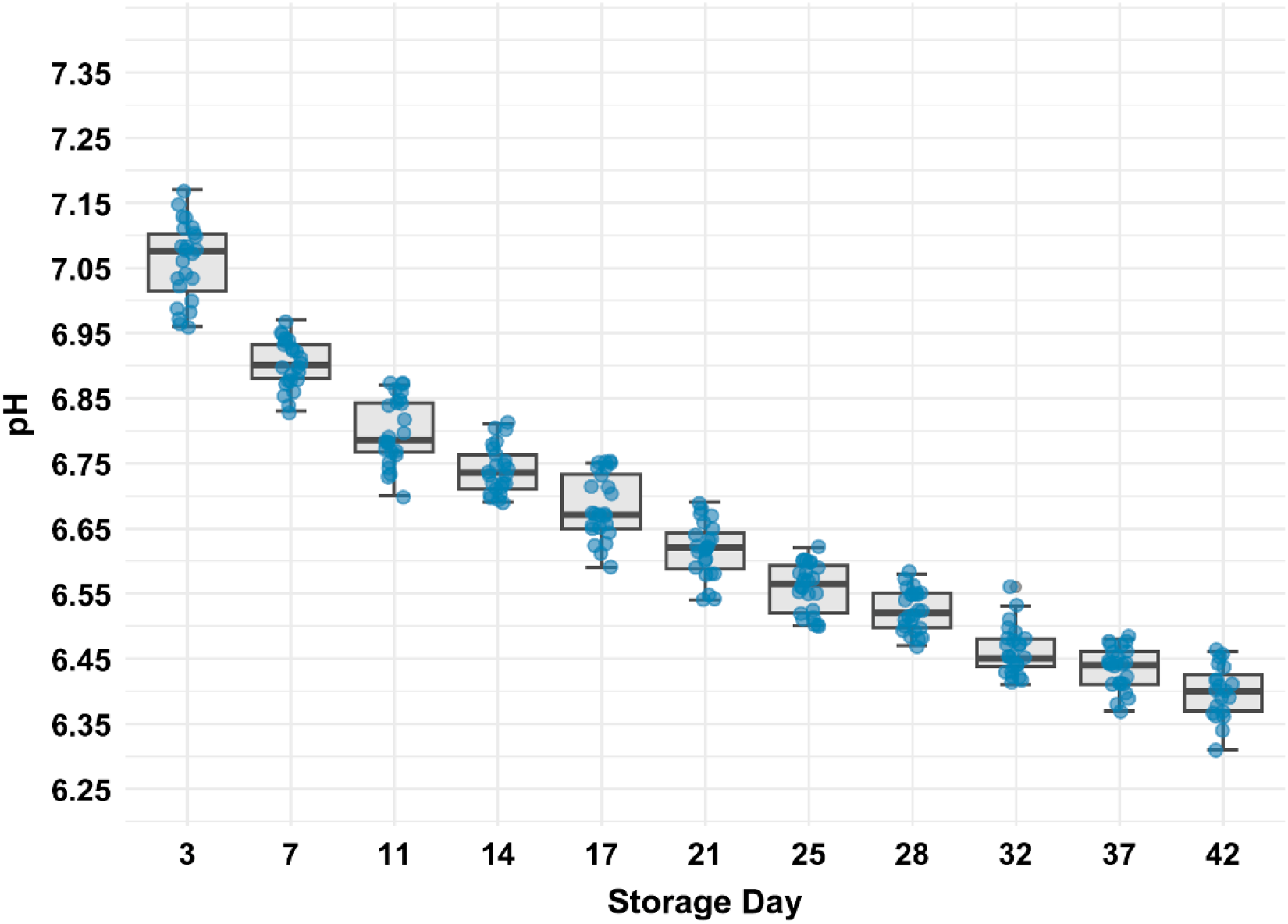
Trajectory of pH values during storage of leukodepleted red blood cell units. Individual measurements (blue points) and corresponding boxplots demonstrate a significant, linear decline in pH from Day 3 to Day 42. A linear mixed-effects model confirmed the association between storage duration and pH (*β* = –0.0163 pH units/day; *p* < 0.001), consistent with progressive metabolic acidification (Table 1).

To quantify the magnitude of acidification at clinically relevant time points, model-based contrasts comparing selected storage days with Day 3 were computed (**Table 1**). pH values were significantly lower at all time points. By Day 7, pH had decreased by Δ = −0.157 (95% CI −0.184 to −0.130; p < 0.0001). Reductions intensified by Day 14 (Δ = −0.320; 95% CI −0.347 to −0.293; p < 0.0001) and Day 21 (Δ = −0.443; 95% CI −0.470 to −0.416; p < 0.0001). By Day 42, units exhibited a pronounced decline (Δ = −0.655; 95% CI −0.684 to −0.627; p < 0.0001), reflecting advanced storage-related acidification. The magnitude of decline increased progressively with storage duration, indicating a monotonic trajectory of acidification.

**Table 1.** Model-based contrasts in pH at selected storage days relative to Day 3. Linear mixed-effects model with bag-level random intercepts. Estimated marginal means (EMMs) with 95% confidence intervals (CI). Negative values indicate lower pH compared with Day 3.

| Storage day<br>Comparison | $\Delta$ pH<br>(estimate) | 95% CI | SE | df | p-value |
| --- | --- | --- | --- | --- | --- |
| **DAY 7 VS DAY 3** | -0.157 | -0.184 to -0.130 | 0.00981 | 226 | <0.0001 |
| **DAY 14 VS DAY 3** | -0.320 | -0.347 to -0.293 | 0.00981 | 226 | <0.0001 |
| **DAY 21 VS DAY 3** | -0.443 | -0.470 to -0.416 | 0.00981 | 226 | <0.0001 |
| **DAY 42 VS DAY 3** | -0.655 | -0.684 to -0.627 | 0.01030 | 227 | <0.0001 |

## Discussion

At the tissue level, O_2_ release from Hb is governed by the Hb–O_2_ dissociation curve, which is strongly modulated by the erythrocyte metabolite 2,3-DPG. The allosteric role of 2,3-DPG in modulating Hb–O_2_ affinity was first established by Benesch and Benesch in 1967 [6]. Subsequent work, including that of MacDonald, further expanded these findings by integrating physiological and clinical perspectives [17]. By binding preferentially to deoxygenated hemoglobin, 2,3-DPG stabilizes the low-affinity tense state, thereby facilitating oxygen unloading to peripheral tissues. Using LC–MS, a reference analytical method, we showed that stored RBCs undergo a rapid, early, and time-dependent decline in 2,3-DPG levels. Notably, this early phase was characterized by substantial inter-unit variability in baseline 2,3-DPG concentrations, detectable as early as Day 3 of storage, before a uniform downward trajectory emerged across all units. The decline was most pronounced during the first week post-collection, culminating in near-complete depletion within two weeks of storage. These results confirm and extend previous reports of progressive 2,3-DPG depletion during RBC storage, with higher-resolution temporal and quantitative characterization than enzymatic assay–based studies [17–19]. The pronounced inter-unit variability in 2,3-DPG levels observed at Day 3 likely reflects heterogeneity present before or immediately after initiation of storage [20]. Donor-specific factors, including differences in glycolytic metabolism and RBC age distribution, influence intracellular 2,3-DPG levels at the time of collection and likely contribute to the broad inter-donor variability observed early in storage [20,21]. As 2,3-DPG levels decline, the Hb-O_2_ dissociation curve shifts leftward, reflecting increased Hb-O_2_ affinity and a reduced capacity for oxygen release at the tissue level [7,11,12].

The mechanisms underlying 2,3-DPG depletion during RBC storage are well established. Stored RBCs rely exclusively on anaerobic glycolysis for energy production and therefore continuously generate lactate within a closed system, without access to the Cori cycle for lactate recycling. Progressive lactate accumulation leads to intracellular acidification, which disrupts glycolytic regulation and directly promotes 2,3-DPG degradation [4,22]. Consistent with these mechanisms, we observed a monotonic decline in storage pH, with mean values reaching 6.40 ± 0.04 by Day 42. This acidification inhibits 2,3-DPG mutase, the enzyme responsible for 2,3-DPG synthesis, and activates 2,3-DPG phosphatase, thereby accelerating 2,3-DPG degradation [17,22].

Based on storage duration, RBC units are frequently labeled as “young” (< 14–21 days of storage) or “old” (> 21 days of storage); however, these designations are largely arbitrary, lack standardized biological justification, and are inconsistently applied across studies, thereby limiting their interpretability and comparability [23,24]. In a meta-analysis including 21 studies— only three of which were randomized—Wang et al. reported that transfusion of older RBC units was associated with a significantly increased risk of mortality (odds ratio 1.16; 95% confidence interval 1.07–1.24) [25]. In contrast, a subsequent meta-analysis by Cheng et al., encompassing 26 randomized controlled trials with low risk of bias, found that transfusion of fresher RBCs did not reduce mortality, transfusion-related adverse events, or length of hospital stay across multiple clinical settings, including critical care and cardiac surgery [24]. Despite the absence of a definitive clinical answer to this question, it is evident that under standard cold-storage conditions, the quality of stored blood deteriorates progressively over time. Reduced 2,3-DPG levels represent only one of several biochemical signatures of RBCs stored for longer than two weeks. This depletion increases hemoglobin’s affinity for oxygen, such that transfused RBCs initially exhibit a diminished capacity to release oxygen to recipient tissues. Although intracellular 2,3-DPG concentrations typically normalize within 24–72 hours after transfusion [7,26], the biological cost of this metabolic restoration—particularly in physiologically vulnerable patients— remains uncertain. Therefore, the apparent lack of clinical benefit of transfusing fresher versus older blood should be interpreted with caution, as RBCs stored for more than two weeks already exhibit substantial biochemical deterioration, and further storage beyond approximately 28 days is associated with increased osmotic fragility and progressive membrane instability [3].

Given the substantial logistical and economic costs associated with modifying RBC transfusion policies, research efforts should prioritize strategies aimed at maintaining intracellular 2,3-DPG levels during storage rather than altering current transfusion practices. Notably, as early as the 1970s, Wood and Beutler demonstrated that supplementation of storage solutions with sodium ascorbate could substantially preserve intracellular 2,3-DPG levels during prolonged refrigerated storage [27].

Subsequent experimental studies have shown that stabilizing storage pH—most notably through removal of CO_2_ using an internal absorbent trap—can partially prevent the decline in intracellular 2,3-DPG levels during RBC storage, thereby attenuating a central metabolic component of the storage lesion [28]. In parallel, biochemical rejuvenation and oxygen-controlled storage strategies have been explored to preserve ATP and 2,3-DPG during prolonged storage. Rejuvenation with metabolic substrates such as pyruvate, inosine, adenine, and phosphate can restore intracellular energy stores late in storage, whereas anaerobic or hypoxic storage limits oxidative and acidotic stress on glycolytic enzymes, stabilizing metabolic flux and preserving 2,3-DPG levels and hemoglobin’s oxygen-unloading capacity [8,12,23,26– 30]. Together, these findings indicate that storage-associated declines in 2,3-DPG represent a potentially modifiable component of the RBC storage lesion. However, whether such approaches can reliably prevent 2,3-DPG depletion or avert storage-related deterioration of RBC morphological phenotypes remains uncertain, underscoring the need for further investigation.

## Conclusions

Using LC–MS as a reference analytical method, we provide a high-resolution reassessment of the timing and extent of 2,3-DPG depletion during RBC storage. Our results demonstrate that key metabolic alterations arise early and reproducibly and are likely driven by the pronounced acidification occurring during the initial storage period. We suggest that 2,3-DPG depletion represents a potentially modifiable component of the RBC storage lesion, supporting future efforts focused on optimizing storage conditions and exploring additive solutions rather than altering transfusion practices based solely on storage duration. Moreover, LC–MS–based metabolomic profiling offers a robust framework for future studies to benchmark metabolic preservation strategies and to integrate biochemical, structural, and functional endpoints when evaluating their potential clinical impact.

## Abbreviations

2,3-DPG: 2,3-diphosphoglycerate
ANOVA: analysis of variance
CPD: citrate-phosphate-dextrose
ESI: electrospray ionization
Hb: hemoglobin
HClO4: perchloric acid
K_2_CO_3_: potassium carbonate
LC-MS: liquid chromatography–mass spectrometry
LMM: linear mixed-effects modeling
OF: osmotic fragility
PBS: phosphate-buffered saline
pH: potential of hydrogen
pRBC: packed red blood cells
RBC: red blood cell
SAGM: Saline, Adenine, Glucose, and Mannitol
SD: standard deviation
WB: whole blood

## Transparency, Rigor and Reproducibility Summary

### Study registration

This study was not formally pre-registered. It was conducted as part of a broader ethically approved research program (protocol ID: PR-AG-411/2017, Ethics Committee of the Vall d’Hebron Research Institute), and the primary objective and analytical approach were defined prior to data collection.

### Analytic plan specification

The analysis plan was not formally pre-registered. However, the team member with primary responsibility for statistical analysis (J. Sahuquillo) certifies that the plan — including the use of linear mixed-effects models (LMMs) with random intercepts for pRBC unit, separate models for storage day and pH as predictors, and Satterthwaite degrees of freedom for fixed-effect inference — was pre-specified before analysis commenced.

### Statistical power and sample size

A sample of 24 leukodepleted pRBC units was planned based on availability of units from the Blood and Tissue Bank and on logistical constraints inherent to the storage protocol. Formal a priori power calculations were not performed, as no published effect size estimates for LC–MS–based 2,3-DPG quantification in SAGM-stored pRBC units were available at the time of study design. The observed effect size for the storage day predictor was β = −0.298 µmol/g Hb per day (robust SE = 0.053; t(18.1) = −5.59; p < 0.001).

### Subject accounting

Twenty-four pRBC units were obtained and included in the study. All 24 units were sampled at each scheduled time point; no units were lost to follow-up or excluded. After Day 17, 2,3-DPG concentrations fell below the LC–MS detection limit in most units; these values were treated as below the limit of quantification and excluded from LMM fitting for that analyte. pH measurements were obtained from all 24 units across all 11 scheduled time points (Days 3–42), with 4 units lost by Day 42.

### Randomization

Randomization was not applicable, as pRBC units were not assigned to intervention groups. Units were processed and sampled sequentially according to the storage timeline.

### Blinding

LC–MS analyses were performed at an external laboratory (Applied Metabolomics Laboratory, IMIM) by investigators not involved in study design or data interpretation, providing de facto analytical blinding. pH measurements were performed without knowledge of LC–MS results at the time of measurement.

### Experimental conditions

pRBC units were maintained at 1–6 °C throughout storage. Samples for LC–MS were collected using heparinized tubes, preprocessed immediately, and stored at −80 °C prior to batched analysis. pH measurements were performed at room temperature (20–23 °C) to reflect conditions at the time of transfusion. All LC–MS analyses were performed in a single batch at IMIM using an ACQUITY UPLC/Xevo TQS Micro system with isotopically labelled internal standards.

### Analytical characteristics

Quantification was performed by external calibration with defined analyte concentrations, complemented by ^13^C-substituted internal standards added to all calibration solutions and sample matrices. The lower limit of quantitation for 2,3-DPG was operationally defined as the minimum detectable concentration above background; samples below this threshold after Day 17 were excluded from LMM analyses.

### Replication

No external replication or validation studies have been performed or are planned at this time.

### Data and code availability

The complete anonymized dataset, metadata, variable dictionary, and R code used for all statistical analyses are available at https://doi.org/10.5281/zenodo.20786075 and at the corresponding GitHub repository: https://github.com/Sahuquillo/dpg-prbc-storage-lesion. Both repositories are linked and cross-referenced.

## Author contributions to the study

Conceptualization: Poca and Sahuquillo; Methodology: Ortuño, Pons, Poca, Sahuquillo, and Peris; Data curation: Peris and Sahuquillo; Investigation: Sahuquillo, Poca, Peris and Rierola; Data analysis and Statistics: Peris and Sahuquillo; Writing – Original Draft: Poca, Sahuquillo and Peris; Writing – Review & Editing: all authors; Funding Acquisition: Sahuquillo, Poca; Resources: Ortuño, Pons, Rierola, Poca and Sahuquillo; Supervision: Sahuquillo and Poca; Role of the Funder/Sponsor: funders had no role in the design and conduct of the study, neither in the collection, analysis, and interpretation of the data not in the review or approval of the manuscript.

## Artificial Intelligence Disclaimer

During the preparation of this manuscript, the authors used Microsoft Copilot and Google Gemini solely for language refinement, stylistic editing, and assistance in debugging R code for statistical analysis. Research Rabbit (https://www.researchrabbit.ai) was used to facilitate literature discovery and reference exploration. No generative AI tools were used for hypothesis development, scientific interpretation, data analysis, or the generation of original scientific content. All study design, statistical analyses, data interpretation, and final conclusions were conceived, executed, and critically reviewed exclusively by the authors, who take full responsibility for the accuracy, integrity, and originality of the work.

